# SkewIT: Skew Index Test for detecting mis-assembled bacterial genomes

**DOI:** 10.1101/2020.02.27.968214

**Authors:** Jennifer Lu, Steven L Salzberg

## Abstract

GC skew is a phenomenon observed in many bacterial genomes, wherein the two replication strands of the same chromosome contain different proportions of guanine and cytosine nucleotides. Here we demonstrate that this phenomenon, which was first discovered in the mid-1990s, can be used today as an analysis tool for the 15,000+ complete bacterial genomes in NCBI’s Refseq library. In order to analyze all 15,000+ genomes, we introduce a new method, SkewIT (Skew Index Test), which calculates a single metric representing the degree of GC skew for a genome. Using this metric, we demonstrate how GC skew patterns are conserved within certain bacterial phyla, e.g. Firmicutes, but show different patterns in other phylogenetic groups such as Actinobacteria. We also discovered that outlier values of SkewIT highlight potential bacterial mis-assemblies. Using our newly defined metric, we identify multiple mis-assembled chromosomal sequences in NCBI’s Refseq library of complete bacterial genomes.

**Software Availability:** SkewIT scripts for analysis of bacterial genomes are provided in the following repository: https://github.com/jenniferlu717/SkewIT.

## Background

Two of the largest and most widely-used nucleotide databases are GenBank [1], which has been a shared repository for more than 25 years (and which is mirrored by the EMBL and DDBJ databases [1, 2]), and RefSeq, a curated subset of GenBank [2]. For sequences to be entered into RefSeq, curators at NCBI perform both automated and manual checks to ensure minimal contamination and high sequence quality. Despite these efforts, multiple studies have identified contamination in RefSeq and other publicly available genome databases [3, 4, 5, 6, 7]. NCBI requires Refseq assemblies to have an appropriate genome length as compared to existing genomes from the same species, and it labels assemblies as “complete” if the genome exists in one contiguous sequence per chromosome, with no unplaced scaffolds and with all chromosomes present. However, NCBI does not perform additional checks, most of which would be computationally expensive, to ensure that a genome sequence was assembled correctly. In this study, we propose a new method, SkewIT (Skew Index Test), for validating bacterial genome assemblies based on the phenomenon of GC-skew. We applied this method to 15,067 complete bacterial genomes in RefSeq, identifying many potential misassemblies as well as trends in GC-skew that are characteristic of some bacterial clades.

### Bacterial GC Skew

GC skew is a non-homogeneous distribution of nucleotides in bacterial DNA strands first discovered in the mid-1990s [8, 9]. Although double-stranded DNA must contain precisely equal numbers of cytosine (C) and guanine (G) bases, the distribution of these nucleotides along a single strand in bacterial chromosomes may be asymmetric. Analysis of many bacterial chromosomes has revealed two distinct compartments, one that is more G-rich and the other that is more C-rich.

Most bacterial genomes are organized into single, circular chromosomes. Replication of the circular chromosomes begins at a single point known as the origin of replication (*ori*) and proceeds bidirectionally until reaching the replication terminus (*ter*). Because the replication process only adds DNA nucleotides to the 3’ end of a DNA strand, it must use two slightly different DNA synthesis methods to allow bidirectional replication of the circular chromosome. The leading strand is synthesized continuously from the 5’ to 3’ end. The lagging strand, in contrast, is synthesized by first creating small Okazaki DNA fragments [10] that are then added to the growing strand in the 3’ to 5’ direction.

These two slightly different replication processes lead to different mutational biases. Notably, the DNA polymerase replicating the leading strand has a higher instance of hydrolytic deamination of the cytosine, resulting in C→ T (thymine) mutations [11]. However, the replication mechanisms for the lagging strand have a higher instance of repair of the same C → T mutation [12]. These differences between the leading and lagging strands result in GC-skew, where the leading strand contains more Gs than Cs, while the lagging strand has more Cs than Gs.

Linear bacterial genomes also exhibit GC skew despite the difference in genome organization. For example, DNA replication of *Borrelia burgdorferi* begins at the center of the linear chromosome and proceeds bidirectionally until reaching the chromosome ends [13, 14]. This bidirectional replication shows the same GC-skew pattern seen on circular chromosomes.

### Quantitative measurements of GC Skew

Since the 1990s, GC skew has been used as a quantitative measure of the guanine and cytosine distribution along a genome sequence, where GC skew is computed using the formula (G-C)/(G+C), where G is the number of guanines and C is the number of cytosines in a fixed-size window [9]. GC skew plots are generated by calculating GC skew in adjacent or overlapping windows across the full length of a bacterial genome [8]. Analysis of these plots confirmed the separation of many bacterial genomes into a leading strand with largely positive GC skew and a lagging strand with negative GC skew. The GC skew effect is strong enough that it can be used to identify, within a few kilobases, the *ori*/*ter* locations.

GC skew plots then evolved into cumulative skew diagrams, which sum the GC skew value in adjacent windows along the bacterial genome [9]. These diagrams sometimes allow more precise identification of the *ori*/*ter* locations, where the origin is located at the global minimum and the terminus is at the global maximum.

### GC Skew Applications and Analyses

Over the last two decades, researchers have employed both GC skew and cumulative GC skew (CGS) diagrams to analyze bacterial genomes. Initial studies confirmed that GC skew was a strong indicator of the direction of replication in the genomes of *Escherichia coli* [15], *Bacillus subtilis, Haemophilus influenzae*, and *Borrelia burgdorferi* [8]. In 1998, Mclean et. al. compared GC-skew among 9 bacterial genomes and 3 archaeal genomes, revealing strong GC-skew in all 9 bacteria but weak or no GC-skew signals in the archaeal genomes [16]. In 2002, Rocha et. al. used CGS to predict ori/ter locations for 15 bacterial genomes [17] and in 2017, Zhang et. al. analyzed GC skew across more than 2000 bacterial genomes [18].

Although GC skew has been used as an indicator of the replication strand in thousands of bacterial genomes, it is rarely used as a means to validate genome assemblies. However, the association between GC skew and replication is strong enough that when a genome has a major mis-assembly such as a translocation or inversion, the GC skew plot is clearly disrupted [19]. We decided to use GC skew to probe the 15,000+ complete bacterial genomes in NCBI’s Refseq library. In order to analyze all 15,000+ genomes efficiently, we introduce SkewIT (Skew Index Test) as an efficient method to calculate the degree of GC skew in a genome. Below we demonstrate how the degree of GC skew tends to be conserved within certain bacterial taxa; e.g. *Klebsiella* species have high values of the skew index, while *Bordetella* have much lower values. During this analysis, we discovered that bacterial genomes with outlier values of SkewIT are highly likely to contain mis-assemblies. Using our newly defined metric, we identify multiple mis-assembled chromosomal sequences in the Refseq library of complete bacterial genomes.

## Method

SkewIT quantifies GC skew patterns by assigning a single value between 0 and 1 to the complete chromosomal sequence of a bacterial genome, where higher values indicate greater GC skew, and lower values indicate that no GC skew pattern was detected. **Figure 1** illustrates the overall method.

**Figure 1.**
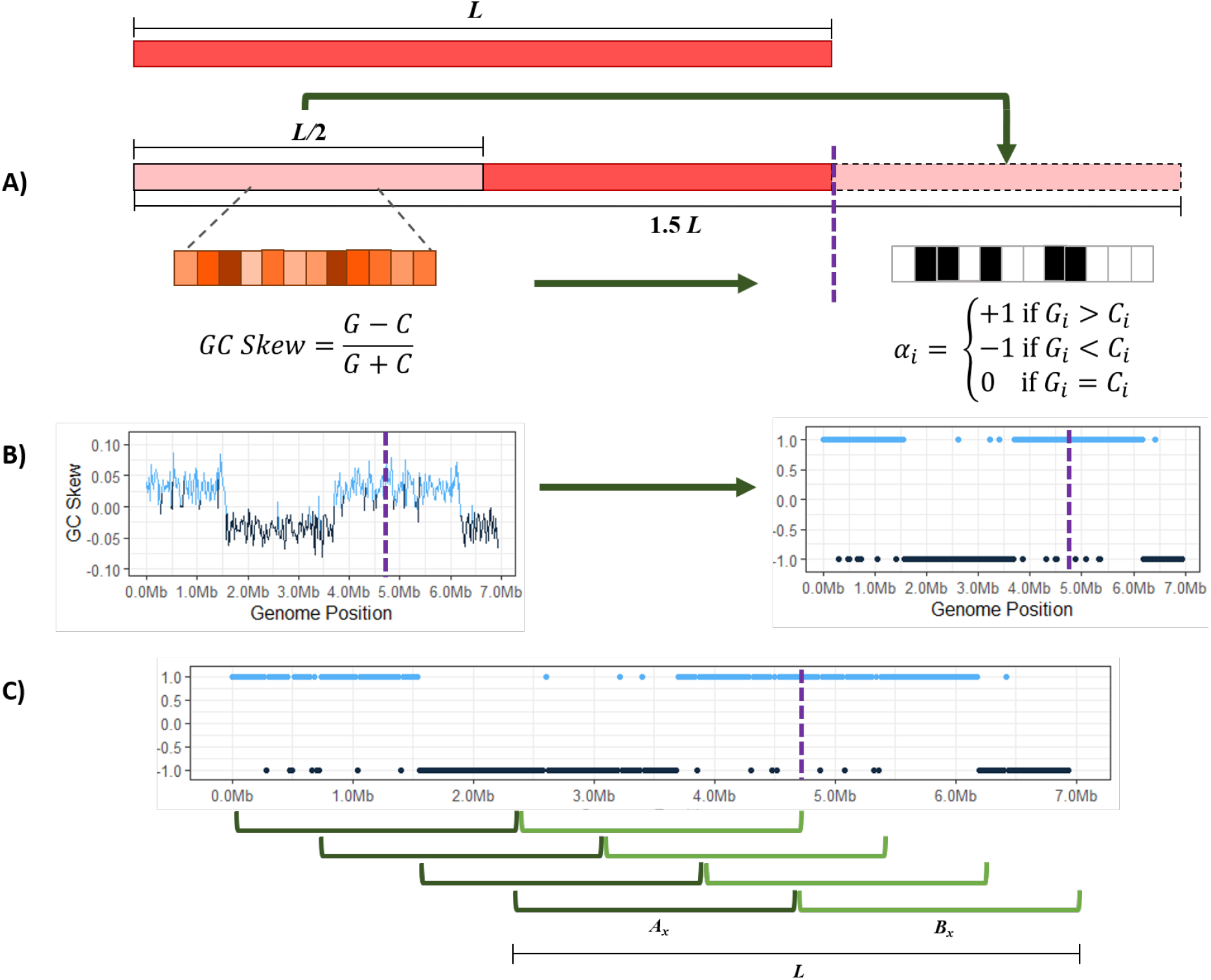
The SkewIT algorithm. A genome of length L is “circularized” by taking the first half of the sequence (L/2) and concatenating that sequence onto the end of the genome (A). The algorithm then splits the sequence into many shorter windows of length *w*. We assign each window an *α* value [1,-1,0] based on whether there are more Gs, Cs, or equal quantities of both. (B) The GC skew statistic is shown (left) plotted across the E. coli genome, with a purple dotted line showing where the original sequence ended, prior to concatenating 1/2 of the genome to the end. The plot on the right shows the *α* value plotted for the same genome. (C) SkewIT finds the location in the genome with the greatest difference in GC skew between the first half and the second half of the genome, by using a pair of sliding windows to find the greatest sum of differences between the *α* values for the two halves.

Although many published bacterial genome assemblies set the origin of replication as the start of the published assembly (i.e., position 1), many other bacterial genomes set coordinate 1 arbitrarily. (Because the genomes are circular, there is no unambiguous choice for the beginning of the sequence. DNA databases only contain linear sequences, and therefore some coordinate must be chosen as position 1.) Therefore, we first “circularize” each bacterial genome of size *L* by appending the first *L*/2 bases of the genome to the end, resulting in a sequence length of 1.5*L* (**Figure 1A**). This ensures that the full genome starting from the origin of replication will be contained within one of the subsequences of length L between position 0 and position L/2.

Next, we select a GC skew window size *w* and split the genome into 1.5*L*/*w* adjacent windows; e.g., for a 1-megabase genome with a 10-Kb window length, we would create 150 windows. In each window *i* ∈ [1,2, ··· 1.5*L*/*w*], we count the frequency of guanine (G) and cytosine (C) bases. Traditionally, GC skew was calculated for each window using **Equation** (1):

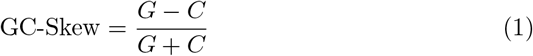

Although the GC skew formula accounts for the relative quantities of G and C bases, our method only evaluates which base is more prominent in each window. **Figure 1B** demonstrates how we convert the GC skew formula into a simplified version that instead assigns each window a score αi using **Equation** (2):

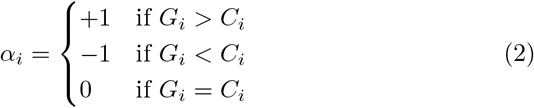

We evaluate the “skewness” of the genome using a sliding window of size *L*, sliding over one window width at a time. For each window *x* ∈ [1, 2, ··· 0.5*L*/*w*], we first sum the *α_i_* values on the left half and separately on the right half of the window (the genome). We calculate the absolute difference between the two sums, as shown in **Equation** (3) and **Figure 1C**:

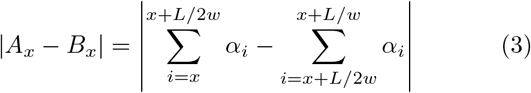

*A_x_* is the sum of the *α* values within the first half of the window, and *B_x_* is the sum of the *α* values for the second half. For example, **Equation** (4) shows how we calculate |*A*_1_ – *B*_1_|, the skewness for the first sliding window from a genome.

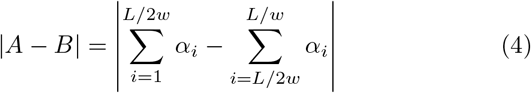

Finally, we determine the maximum value of |*A_x_* – *B_x_*|, which gives us the window where the greatest difference exists between the GC content of one half of the genome and the GC content of the other half. In order to be provide a consistent value between 0 and 1 despite genome length *L* or window size *w*, we define the skew index (*SkewI*) as the following normalized value:

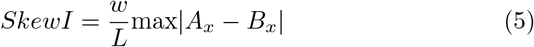

## Results and Discussion

We applied the SkewIT method using a window size of 20Kb to the complete bacterial genomes from NCBI RefSeq Release 97 (released on November 4, 2019). We only evaluated bacterial chromosomes that were > 50,000bp in length and excluded plasmids from this analysis. In total, we tested 15,067 genomes representing 4,471 species and 1,148 genera. **Supplemental Table 1** lists each genome with their SkewI values. The table also provides the main taxonomy assignments for each genome.

Overall, analysis of all bacteria revealed that most genomes have strong GC skew patterns, with relatively few having SkewI values less than 0.5 (**Supplemental Figure 1**). In order to isolate and analyze bacterial genomes with unusually low SkewI values, we separated the bacterial genomes by clades, revealing characteristic SkewI distributions for individual genera (**Figure 2**). For example, genomes from the genera of *Bacillus, Escherichia*, and *Salmonella* have consistently high SkewI values, with a mean close to 0.9. However, *Bordetella* genomes have far lower SkewI values, with a mean of 0.45. Additionally, while genomes in the *Klebsiella* and *Brucella* genera all have similar SkewI values (and therefore similar amounts of GC skew), genomes from the *Campylobacter* and *Corynebacterium* genera demonstrated much less consistent amounts of GC skew, with a wide range of SkewI values.

**Figure 2.**
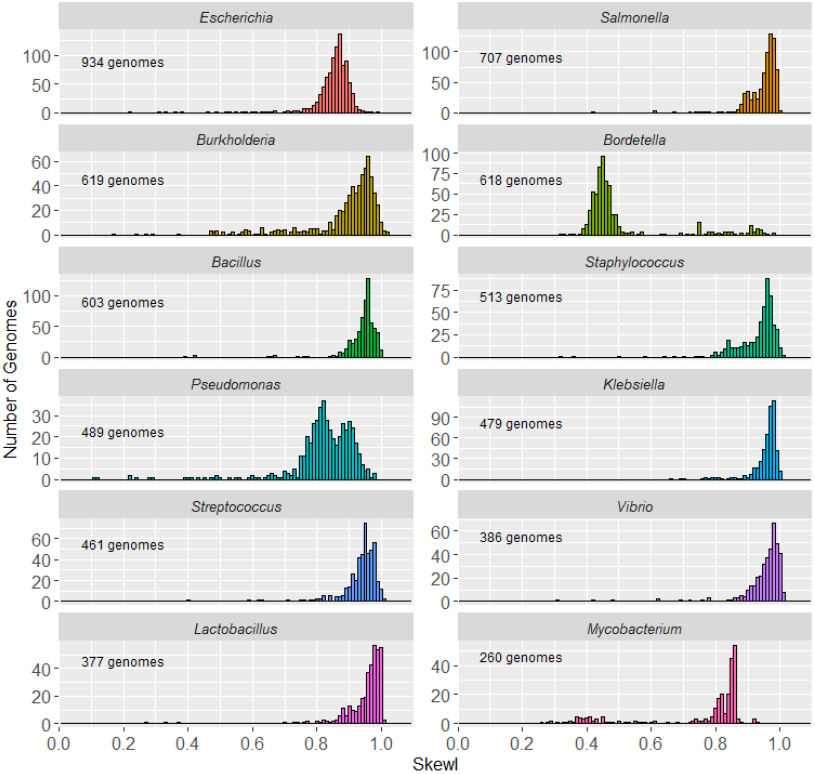
Skew index (SkewI) per genus. This figure shows the distribution of SkewI values for the 12 bacterial genera with the greatest number of fully sequenced genomes.

Given the differences between genera, we evaluated abnormalities in GC skew by setting a threshold for each genus that would allow us to flag genomes that might have assembly problems. For each genus with 10 or more genomes, we set a SkewI threshold at two standard deviations below the mean (**Table 1** and **Supplemental Table 2**). If a genome’s SkewI exceeded the threshold, then we considered that bacterial genome to be within the expected range for that genera. However, if a genome’s SkewI was below the threshold, then we considered that genome to be possibly mis-assembled.

**Table 1.** Average SkewI values for the 12 bacterial genera with the largest number of complete genomes. The threshold was set at 2 standard deviations below the mean.

| Genus | Genome Count | Mean | st. dev. | Threshold |
| --- | --- | --- | --- | --- |
| Escherichia | 934 | 0.8486 | 0.0688 | 0.7110 |
| Salmonella | 707 | 0.9482 | 0.0502 | 0.8478 |
| Burkholderia | 619 | 0.8926 | 0.1181 | 0.6564 |
| Bordetella | 618 | 0.4988 | 0.1394 | 0.2200 |
| Bacillus | 603 | 0.9422 | 0.0545 | 0.8332 |
| Staphylococcus | 513 | 0.9291 | 0.0697 | 0.7897 |
| Pseudomonas | 489 | 0.8167 | 0.1148 | 0.5871 |
| Klebsiella | 479 | 0.9567 | 0.0430 | 0.8707 |
| Streptococcus | 461 | 0.9373 | 0.0557 | 0.8259 |
| Vibrio | 386 | 0.9526 | 0.0686 | 0.8154 |
| Lactobacillus | 377 | 0.9499 | 0.0748 | 0.8003 |
| Mycobacterium | 260 | 0.7460 | 0.1751 | 0.3959 |
| Acinetobacter | 252 | 0.9445 | 0.0759 | 0.7928 |
| Campylobacter | 248 | 0.7425 | 0.0968 | 0.5489 |
| Corynebacterium | 224 | 0.7890 | 0.2085 | 0.3720 |

During our investigation into each genome with a SkewI below the threshold, we identified several potentially mis-assembled *Escherichia* and *Burkholderia* genomes. Additionally, we were able to identify an interesting phenomenon in *Mycobacterium* genomes relating GC-Skew to GC-content. The following sections describes these findings.

### Escherichia

For the Escherichia genus, RefSeq contains 934 complete genomes, with an average SkewI value of 0.85 and a threshold of 0.71 (**Figure 3A**). While the majority of *Escherichia* genomes had SkewI values above the threshold, one of them, *Escherichia coli O121 strain RM8352* (*E. coli O121*), had a SkewI of 0.223, which appeared far too low. In an effort to validate this assembly, we aligned the original raw reads back to the genome while also comparing *E. coli O121* to *Escherichia coli M8*, which has a more-typical SkewI of 0.869. Initial analysis of the GC-skew plots for both *E. coli* genomes revealed a clear difference between the genomes, as shown in **Figure 3B**. For *E. coli M8*, the GC skew plot shows that almost precisely half the genome has more Gs than Cs, and the other half has more Cs than Gs, as is typical for this species. In *E. coli O121*, by contrast, a much larger portion of the forward strand has more Gs than Cs. We then aligned *E. coli O121* against *E. coli M8* (using used NUCmer [20]), revealing a large inversion in *E. coli O121* from position 2,583,081 to 4,963,263. Alignment of assembly reads to each genome using Bowtie2 [21] revealed gaps in coverage at the points flanking both ends of the inversion in *E. coli O121*, suggested that the assembly is incorrect in those regions (**Figure 3C**).

**Figure 3.**
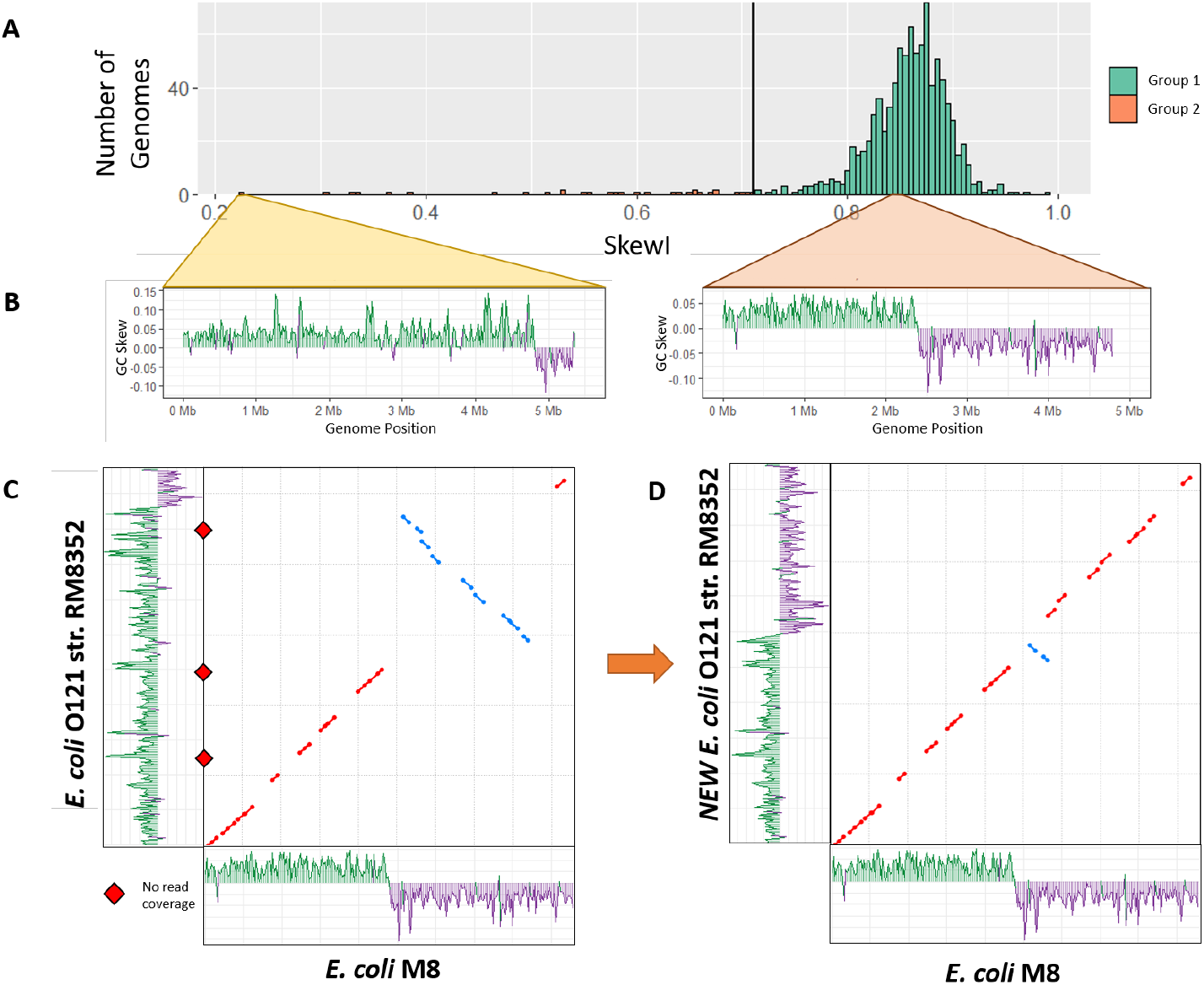
Escherichia skew index values. A) SkewI for all 934 *Escherichia* genomes. The threshold (vertical black line) is at 0.711. B) GC-skew plots for *Escherichia coli O121 strain RM8352* and *Escherichia coli M8. E. coli O121* has an unusually low SkewI of 0.223, while *E. coli M8* has a SkewI of 0.86, which is typical for this genus. C) Initial alignment between the two *E. coli* genomes revealed a large inversion. Alignment of the assembly reads revealed locations with no read coverage (red diamonds) *E. coli O121* at both ends of the inversion. D) Flipping the inversion in strain RM8352 produced a much more consistent alignment between the *E. coli* genomes (dot plot), and restored the GC skew plot to a more normal appearance (shown along the y axis).

Because there were no reads supporting the inversion from 2,583,081 to 4,963,263 in *E. coli O121*, we replaced this sequence with its reverse complement and repeated our analysis. Our new *E. coli O121* genome has a SkewI of 0.77 with an evenly divided GC-skew plot (**Figure 3D**). Comparison of the new *E. coli O121* against *E. coli M8* shows a much more consistent 1-to-1 alignment between the two genomes, with only one small inversion remaining.

### Burkholderia

The *Burkholderia* genomes have a mean SkewI of 0.849 with a SkewI threshold of 0.656 (**Figure 4A**). Although there are 619 finished chromosomes from the *Burkholderia* genus, they represent only 270 individual organisms; each *Burkholderia* strain typically has 2-3 chromosomes. **Figure 4B** shows the SkewI distribution based on chromosome. There is no significant difference in SkewI between chromosomes.

**Figure 4.**
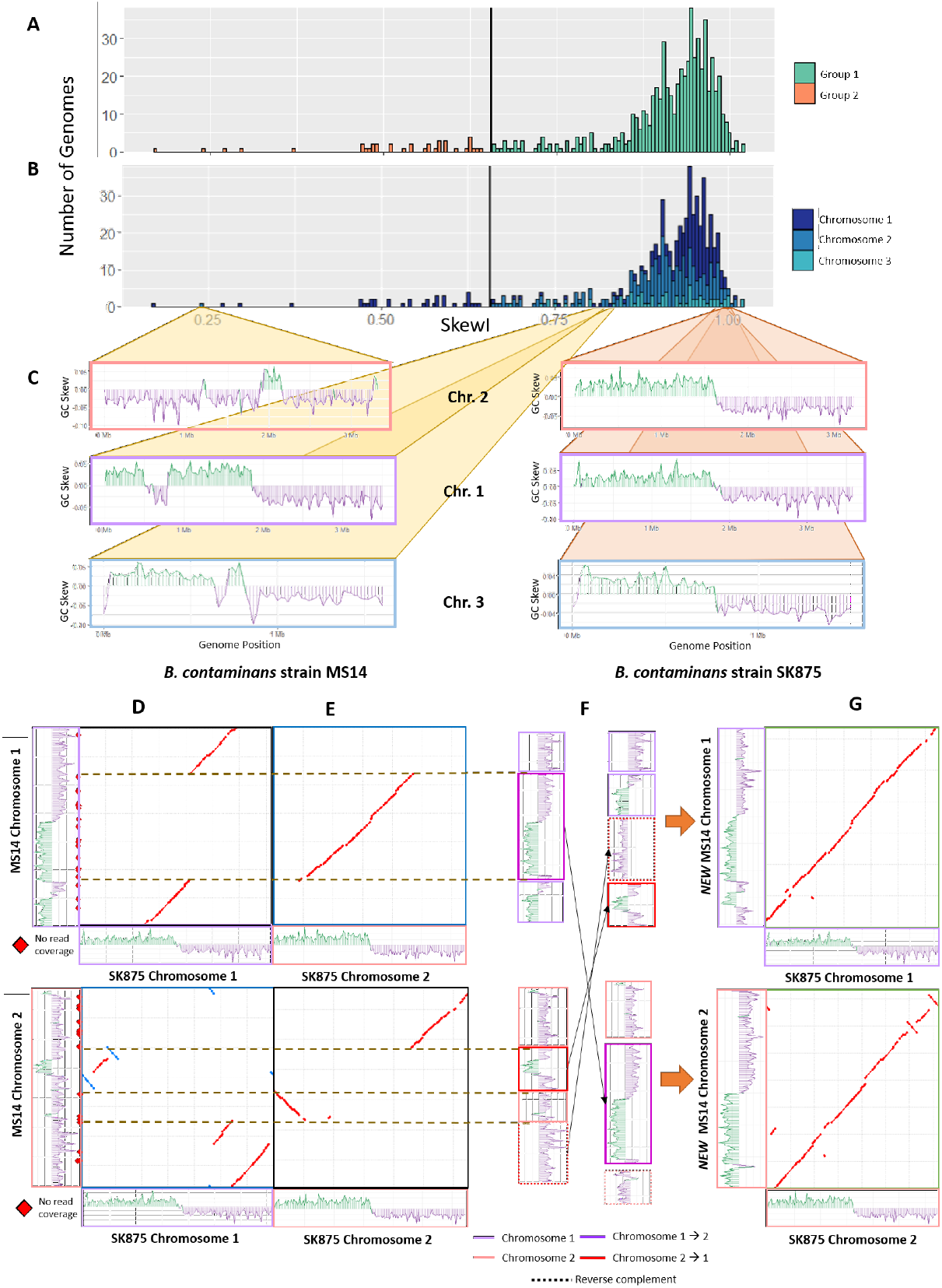
Burkholderia skew index values. A) SkewI for all 934 Burkholderia genomes. The threshold (vertical black line) is 0.656. B) SkewI colored by chromosome. C) GC-skew plots for all three chromosomes for Burkholderia contaminans strains MS14 (left) and SK875 (right). D) Alignments between MS14 and SK875 chromosomes 1 and 2. MS14 is shown on the y axis of each plot. E) Cross-chromosome alignments between MS14 and SK875 chromosome 1 and 2 reveal that a 1.7Mbp region of MS14 chromosome 1 actually belongs to chromosome 2. Similar matches in MS14 chromosome 2 suggest two regions that belong in chromosome 1. F) We rearranged and inverted the sequences of MS14 chromosomes 1 and 2 based on the alignments and GC-Skew plots. G) The final MS14 chromosomes alignment with those of *B. contaminans* SK875.

Further analysis of the individual genomes with SkewI values below the threshold revealed significant differences between the SkewI values for the three chromosomes of *Burkholderia contaminans MS14*. Notably, chromosome 2 had a SkewI of 0.238 while chromosomes 1 and 3 had SkewIs of 0.806 and 0.860 respectively (**Figure 4C**). By comparison, the three chromosomes of a different strain, *Burkholderia contaminans SK875*, all had very high SkewIs of 0.973, 0.997, and 0.974.

Aligning the raw *B. contaminans MS14* assembly reads against the three chromosomes using Bowtie2 [21] revealed many locations with no read coverage, suggesting that the full read set used for the assembly was not available. We then aligned the *B. contaminans MS14* chromosomes against the same chromosomes for *B. contaminans SK875* and observed multiple large-scale disagreements between the chromosomes. While chromosome 3 from both strains aligned nearly perfectly, only 50% of chromosome 1 and 2 of MS14 aligned to the same corresponding chromosome of *B. contaminans SK875* (**Figure 4D**).

We then aligned chromosome 1 of *B. contaminans MS14* to chromosome 2 of *B. contaminans SK875* and vice versa and discovered that the sequences of *B. contaminans MS14* appeared mis-assembled (**Figure 4E**). Based on the differences in alignment and the GC Skew plots of *B. contaminans MS14*, it appears that the 1.7Mbp region of *B. contaminans MS14* chromosome 1 from 812,522 to 2,579,632 belongs to chromosome 2. Similarly, two regions from *B. contaminans MS14* chromosome 2 belong to chromosome 1.

Based on the chromosome alignments and GC-skew plots, we rearranged and inverted the individual *B. contaminans MS14* sequences as illustrated in **Figure 4F**. The final SkewI for these corrected chromosome 1 and chromosome 2 sequences were 0.774 and 0.946 respectively, both within the expected range. Additionally, realigning the new MS14 sequences against those of SK875 a far higher degree of synteny between the two genomes (**Figure 4G**).

### Mycobacterium

Analysis of the *Mycobacterium* SkewI distribution revealed a main peak at 0.85 and a smaller peak centered around 0.4 (**Figure 5A**). Due to the large standard deviation, the SkewI threshold was calculated to be 0.396, with 20 genomes falling below the threshold. However, upon investigation into the individual genomes, it appeared that all 20 of these genomes come from *Mycobacterium avium* and *M. avium* subspecies, suggesting that the SkewI values are not reflective of a mis-assembly but rather reflective of a different degree of skew in *M. avium* and possibly other species within the Mycobacteria.

**Figure 5.**
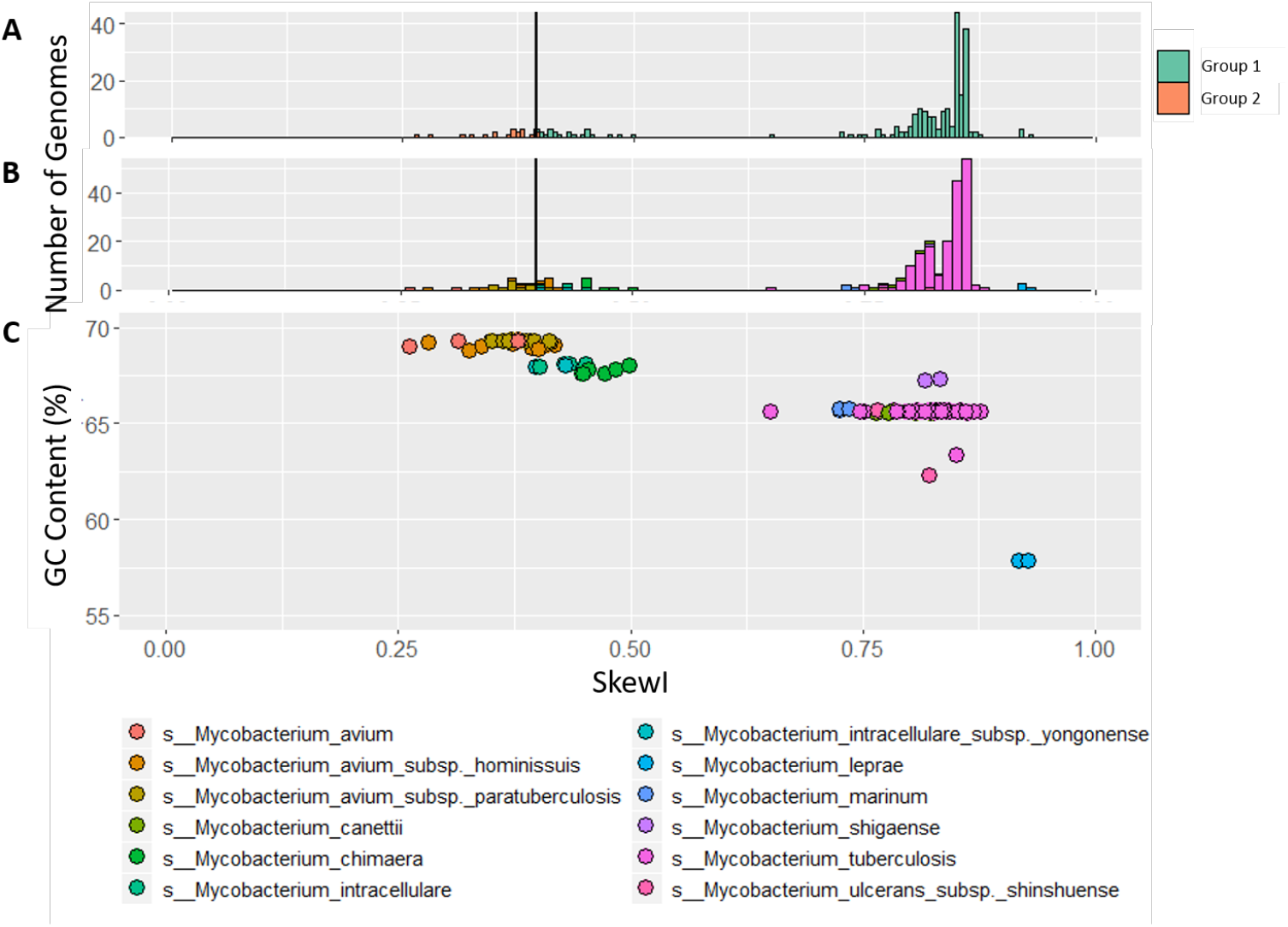
Mycobacterium skew index values. A) SkewI for 236 *Mycobacterium* genomes from 12 *Mycobacterium* species, all of which have multiple strains available in RefSeq. The threshold (vertical line) is at 0.396. B) SkewI colored by species. C) Plot comparing GC Content (%) to SkewI, where each dot represents a different genome colored by species.

We explored this hypothesis by re-plotting SkewI using different colors for each of the 12 species, as shown in **Figure 5B**. As the plot shows, the large peak centered around 0.85 mainly consists of the 179 *M. tuberculosis* genomes while the smaller peak mainly consists of the 27 *M. avium* genomes. Because *Mycobacterium* genomes have a high GC-content (%), we then plotted GC-content vs. SkewI for these same genomes (**Figure 5C**), revealing that for the *Mycobacterium* genus, higher GC-content results in a lower SkewI.

## Conclusion

Our SkewIT (Skew Index Test) provides a fast method for identifying potentially mis-assembled genomes based on the well-known GC skew phenomenon for bacterial genomes. In this study, we describe an algorithm that computes a new GC-skew statistic, SkewI, and we computed this statistic across the 15,067 genomes from RefSeq, discovering that GC skew varies considerably across genera. We also used anomalous values of SkewI to identify likely mis-assemblies in *Escherichia coli O121 strain RM8352* and in two chromosomes of *Burkholderia contaminans MS14*. We suggest that researchers can validate future bacterial genome assemblies by running SkewIT and comparing the resulting SkewI value to the thresholds in **Supplemental Table 2**. Genomes with SkewI values lower than the expected threshold should be further validated by comparison to closely-related genomes and by alignment of the original reads to the genome.

## Availability of data and materials

The SkewIT github repository (https://github.com/jenniferlu717/SkewIT) provides SkewIT scripts for analysis of bacterial genomes and files containing SkewI values and thresholds for RefSeq Release 97 bacterial genomes.

## Author’s contributions

JL conceived and designed the experiments, performed the experiments, analyzed the data, wrote the paper, prepared figures and/or tables, performed the computation work, and wrote the paper.

SLS conceived and designed the experiments, analyzed the data, and wrote the paper.

## Acknowledgements

We would like to thank Martin Steinegger for helpful comments and feedback on the paper draft and figures. This work was supported in part by NIH under grants R01-HG006677 and R35-GM130151 and by NSF under grant IOS-1744309.

## Supplemental Figures

**Supplemental Figure 1.**
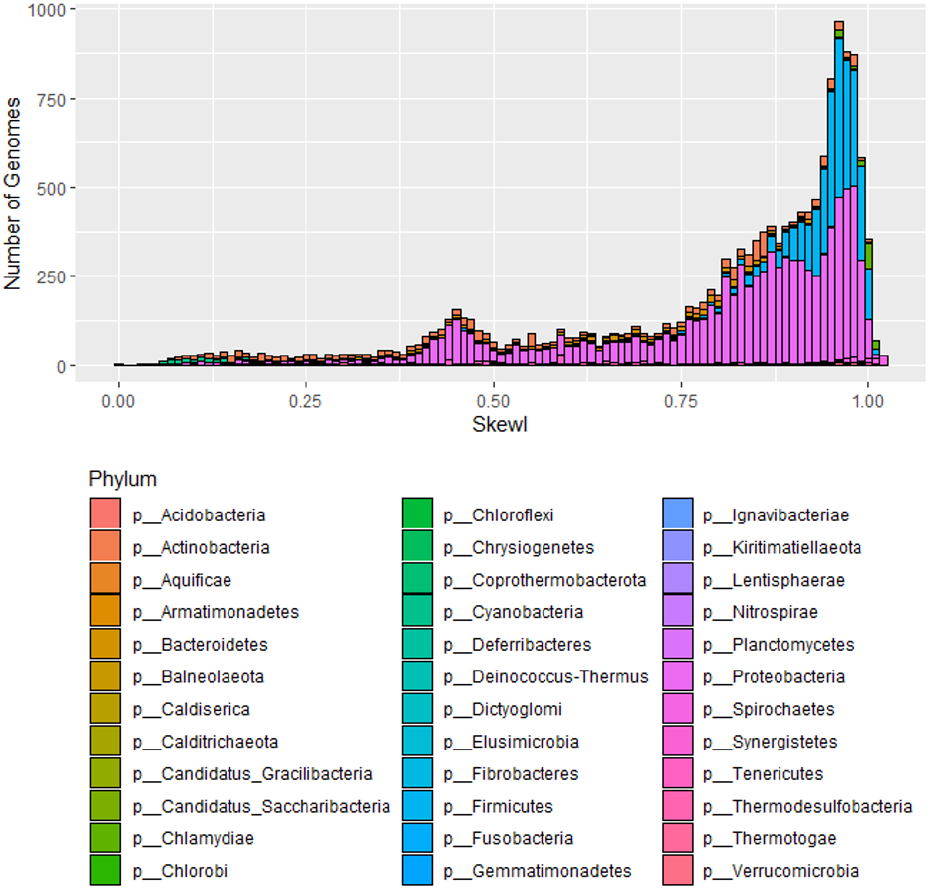
SkewI for all 15,067 complete bacterial RefSeq genomes.

## Supplemental Files

**Supplemental Table 1: Bacterial Genomes SkewI**

All 15,067 bacterial genomes are listed along with their calculated SkewI values. Additionally, this table lists the kingdom, phyla, class, order, family, genus, and species names/NCBI taxonomy IDs for each genome.

**Supplemental Table 2: SkewI Thresholds per genus**

For all bacterial genera analyzed, we list the number of genomes, the average SkewI, and the SkewI standard deviation. For any genus with more than 10 genomes, we also include a threshold, which is 2 standard deviations below the mean.

